# Stage-specific gene ratios highlight genes and mechanisms related to presymptomatic and symptomatic Multiple Myeloma

**DOI:** 10.1101/2024.11.04.621824

**Authors:** Grigoris Georgiou, George Minadakis, Nestoras Karathanasis, Kyriaki Savva, Efi Athieniti, Marilena M Bourdakou, George M Spyrou

## Abstract

**Background/aim:** Multiple Myeloma is the second most common blood cancer, characterised by the accumulation of malignant plasma cells and the production of large amounts of a monoclonal immunoglobulin protein, in the bone marrow. The identification and progression/behaviour of molecular markers across stages remains a scientific challenge. This work aims to provide a holistic approach to the understanding of the disease progression, providing specific methodologies and candidate biomarkers, able to characterise and distinguish the disease state across stages.

**Materials and methods:** Two large bulk RNA datasets were used to collect and integrate stage-specific information at the gene level by means of: (a) differential expression analysis to obtain differential expressed genes (DEGs), (b) a recently introduced computational methodology able to detect monotonically expressed genes (MEGs), (c) a proposed computational methodology that uses pairs of MEGs at sample level, to classify and discriminate different stages of Multiple Myeloma. Additional numerical metrics were applied to rank the performance of these pairs across samples, facilitating the characterization and differentiation of disease stages. Validation was conducted using five additional external datasets, which were then utilised to enrich the final selection of top-rated genes identified from the two bulk RNA datasets under study. The final top-ranked genes were further used for pathway enrichment analysis in order to provide candidate pathways per stage.

**Results:** We first show that MEGs provide better statistics than DEGs, both at gene and pathway level analysis. Secondly, we show that the proposed computational methodology by means of MEGs reveals short lists of high discriminative genes across stages, which in turn highlight significant groups of pathways.

**Conclusion:** We integrated traditional analysis of DEGs with a recently introduced methodology for identifying MEGs, creating a novel computational approach capable of identifying highly discriminative genes and pathways that can serve as candidate markers for stage identification in a single sample.

**Highlights:**

- A novel computational approach was used to identify Monotonically Expressed Genes (MEGs) whose expression was constantly increasing or decreasing. Genes such as RB1, CD27, TP53, and MCL1, previously highlighted in Multiple Myeloma, showed a consistent monotonic pattern, providing potential indicators for tracking the progression from the pre-malignant stages to active Multiple Myeloma.
- The study calculated gene pair ratios using MEGs characterised by normal distribution and low dispersion. These ratios effectively distinguished healthy from disease samples, although the discrimination between disease stages (MGUS, SMM, MM) was less clear due to their overlapping molecular profiles.
- Enrichment analysis of significant gene pairs identified critical pathways affected during Multiple Myeloma progression, such as bone disease-related calcium pathways, glucocorticoid-regulated functions, and cardiac and neurological systems. These findings align with known clinical manifestations in MM patients, such as bone disease and amyloid cardiomyopathy.
- The gene lists generated from our computational approach were validated against internal and external datasets, confirming their applicability. The methodology showed promise in identifying candidate genes for disease progression and could be applied to other diseases to uncover novel gene pairs not highlighted by traditional analyses.

## Introduction

Multiple Myeloma (MM) is the second most common blood cancer, characterised by the accumulation of malignant plasma cells and the production of large amounts of a monoclonal immunoglobulin protein, in the bone marrow [1]. Monoclonal Gammopathy of Undetermined Significance (MGUS) and Smoldering Multiple Myeloma (SMM) are the asymptomatic precursory stages of MM. However, the identification and progression/behaviour of molecular markers across stages remain a scientific challenge. This work aims to provide a holistic approach to the understanding of the disease progression, providing specific methodologies and candidate biomarkers, able to characterise and distinguish the disease state across stages. Recently Bourdakou, et al. (2021), introduced a computational methodology performed on colorectal cancer, based on the concept of Monotonically Expressed Genes (MEGs) and Monotonically Enriched Pathways (MEPs), providing noteworthy results [2]. According to the authors, having the Differential Expression Analysis (DEA) of a stage-specific dataset, MEGs are actually genes with a constant increase or decrease of the log_2_FC parameter across stages, and in turn, are considered to follow a monotonic trend. The underlying approach draws from the assumption that genes with monotonic tendencies may be more related to the progression of the disease in contrast to those that do not have this behaviour. It is a natural characteristic that indicates the direction of change in differential gene expression, which can be particularly informative for understanding disease progression. In this line, we used bulk RNA datasets from patients with Multiple Myeloma, in order to collect and integrate stage-specific information from DEGs and MEGs at gene and pathway levels. First, we examined the statistical significance of MEGs, providing a comparative analysis of their behaviour versus DEGs. In order to examine and advance the use of MEGs towards identifying stages of Multiple Myeloma, we further developed a computational methodology that incorporates numerical parameters to enhance the characterization and differentiation of disease stages. By integrating traditional analysis of DEGs with MEGs, we provide a novel computational approach capable of identifying highly discriminative genes and pathways that can serve as candidate markers for stage identification in a single sample. To ensure the reliability and generalizability of our methodologies, we validated our findings using four external datasets, demonstrating the robustness of our approach and its applicability to independent patient cohorts. This work not only advances the scientific understanding providing a deeper insight into the molecular mechanisms and genetic pathways involved in Multiple Myeloma (MM), but also offers a promising methodology for implementing appropriate tools to improve disease diagnosis, monitoring and treatment.

## Materials and Methods

The workflow in this work can be described by means of the following steps depicted in Figure 1. Specifically, the process begins with Differential Expression Analysis on the selected bulk RNA datasets to identify genes whose expression changes across disease stages. The next step involves applying a recently introduced methodology for identifying genes that exhibit a monotonic trend to their expression as the disease progresses. Combining the two aforementioned approaches, we further introduce a novel computational methodology where ratio distributions of MEGs-specific gene-pairs within each sample, are used for distinguishing between different conditions under study. To enhance the accuracy of this methodology, only the distributions that exhibit normal distribution and small dispersion around their mean were retained for the analysis. The retained gene-pair distributions were further enriched and validated using external datasets, employing classification performance metrics such as Accuracy (ACC) and Area Under the Curve (AUC) which in turn were used to assess the method’s effectiveness. Based on different performance thresholds, lists of significant genes and pathways were further generated and evaluated from a biological perspective. The following sections provide a detailed description of the methodologies used in this study.

**Figure 1:**
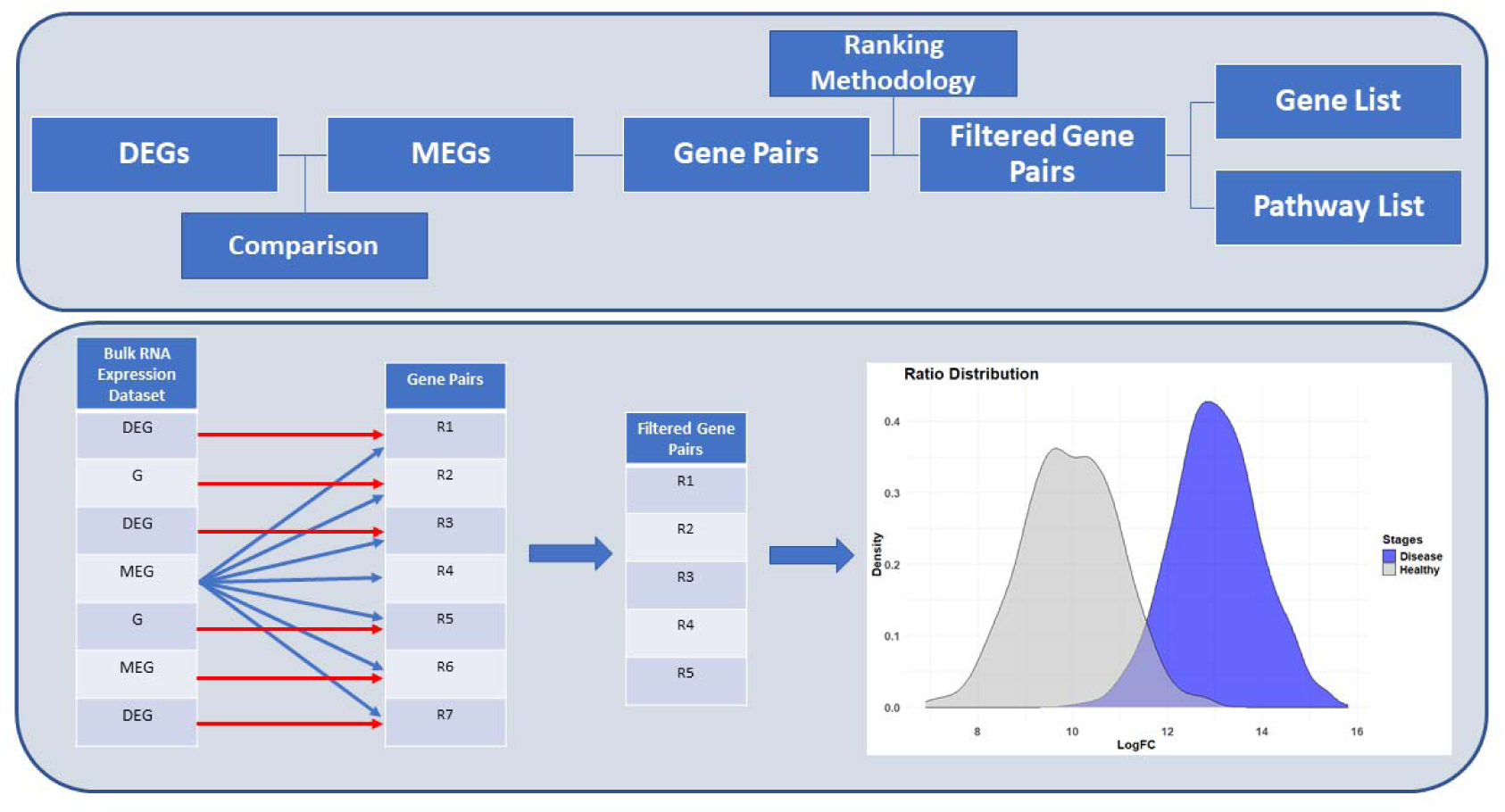
Overview of the workflow followed in this work. Starting with DEA we compute the DEGs between Healthy and Disease stages. Then, we extract those with Monotonic trends in their Log_2_FC based on a previously published methodology. Gene pairs were calculated according to a new computational approach, where Genes (G), DEGs and MEGs are used. Those that followed a normal distribution and small dispersion around their mean were kept forward to make pairwise comparisons between all the available stages. Finally based on the applied metrics used to calculate the performance of our approach, gene and pathway lists were generated.

### Data collection and preprocessing

The data under study have been collected from the Gene Expression Omnibus (GEO) [3], using the GEOquery R package [4]. Specifically, 891 samples were totally collected from seven bulk RNA microarray datasets namely the GSE13591 [5], GSE14230 [6], GSE235356 [7], GSE47552 [8], GSE5900 [9], GSE6477 [10] and GSE80608 [11], which include two or more generic (MGUS, SMM, and MM) stages of multiple myeloma, as shown in the following table.

Herein it should be noted that the main analysis in this work will be focused on the two datasets which contain all the three stages of multiple myeloma, namely the GSE47552 and the GSE6477, accordingly. The rest of the datasets will be further used for testing and evaluation of the results.

### Generating gene signatures related to the progression of the disease

To derive a gene signature and a computational method that exploits this signature towards the identification of stages in Multiple Myeloma, we initially explored the gene signatures derived through (a) differential expression analysis, and (b) monotonic differential expression analysis. Then, we evaluated the benefits of using monotonic differential expression analysis and we developed a classification approach based only on within-sample calculations of gene expression ratios from the monotonic differential expression analysis. This approach aims to create a reference dataset containing highly discriminative MEG ratios across stages, which could potentially be used to identify the stage at the level of a single sample without relying on RNA-Seq dataset comparisons.

### Differential Expression Analysis of Genes

Differential Expression Analysis (DEA) was performed by means of the Limma (version 3.56.2) R package [12], which provides a linear model that calculates a moderated t-statistic from gene expression experiments. Specifically, each dataset was preprocessed utilising quantile normalisation, background correction and log_2_ transformation. Then for each stage comparison, we obtained the fold change (FC) and log_2_FC parameters for each gene, along with the adjusted p-values using the empirical Bayes moderated t-statistics [13]. Figure 2 depicts the number of over- and under-expressed genes across stages, obtained from the two aforementioned datasets under study, showing a consistent pattern of balance across stages.

**Figure 2:**
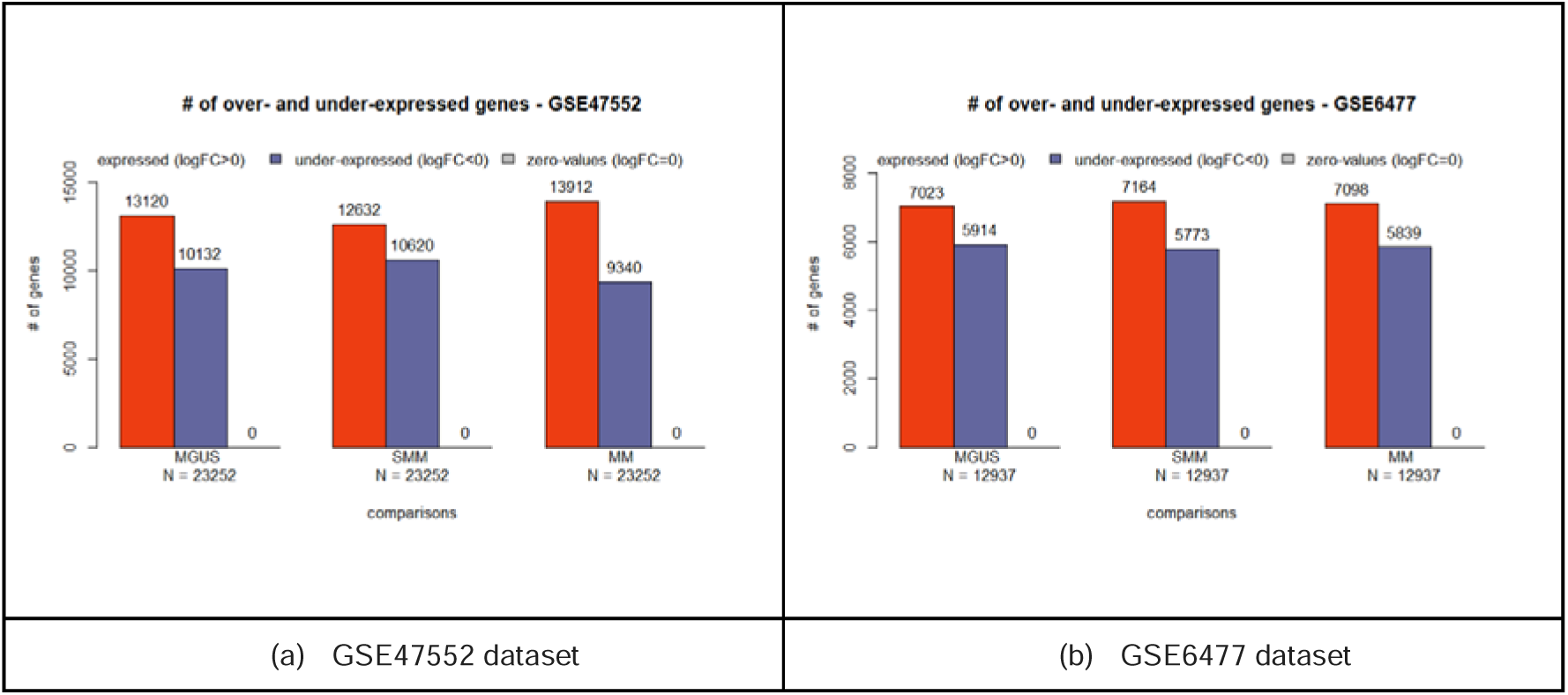
Over- and under-expressed information for the two datasets that include the three stages of multiple myeloma. The red bars refer to the number of over-expressed genes, the blue ones refer to the under-expressed ones, while the grey ones refer to the number of those with zero log_2_FC value.

In order to show the statistical coverage of the obtained adjusted p-value, a specific algorithm was developed to calculate the remaining number of genes for different thresholds of *p_adj_*. This in effect allows the selection of an optional *p_adj_*-threshold that could potentially be used for achieving an optimal filtering of the differentially expressed data. Figures 3a,3b depict the number of remaining genes for different thresholds of *p_adj_*-value for each stage included in the two datasets. It is observed that for the theoretical values of *p_adj_* ≤ 0.05, the remaining number of significant genes range between 2000 and 4000 across stages, which is an acceptable amount. The results from DEA are depicted in the Supplementary Files1-2.

**Figure 3:**
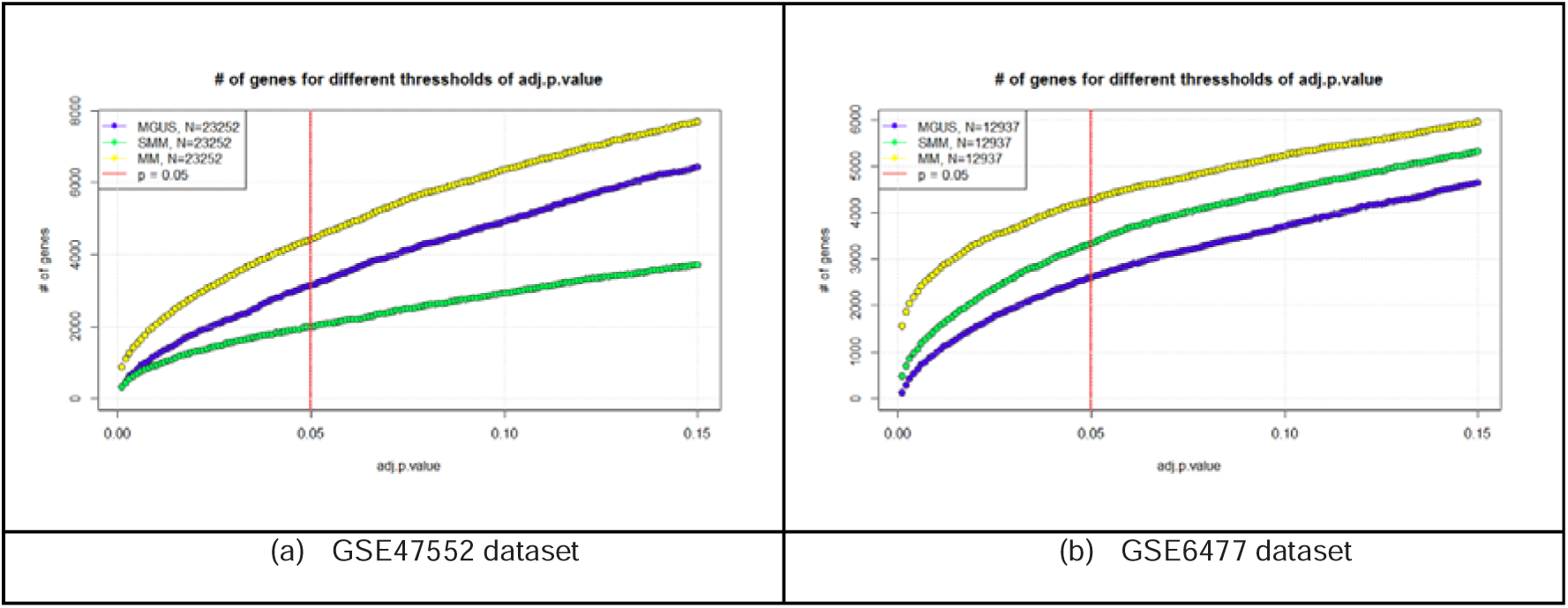
Number of remaining genes for different thresholds of *p_adj_*-value for each stage included in the two datasets. The red vertical line refers to the theoretical value of *p_adj_* < 0.05, normally used for filtering.

### Monotonic Differential Expression Analysis

The term Monotonically Expressed Genes (MEGs) derives from a recently introduced computational methodology performed on colorectal cancer [2] with notable results. According to the authors, having the DEA outcome from a stage-specific dataset, MEGs are actually genes with a constantly increasing or decreasing log_2_FC value across stages, and in turn are considered to follow a monotonic trend. The underlying approach draws from the assumption that genes with monotonic trend may be more related to the disease progression in contrast to those that do not have this behaviour. A necessary condition for selecting datasets in finding monotonic genes is to contain samples from at least three disease stages. Thus, the genes examined for monotonicity were those involved in the two datasets which include all the three stages (MGUS, SMM and MM), namely the GSE6477 and the GSE47552 respectively. Herein, for each stage comparison against healthy samples, we applied the theoretical threshold of *p_adj_* ≤ 0.05, in the prospect to keep only the significant genes included in the two datasets under study (named *original-gene-set*). In consequence using the same threshold of *p_adj_* ≤ 0.05, we further filtered the monotonic genes included in the DEA datasets (named *monotonic-gene-set*). Figure 4 depicts the number of over- and under-expressed genes of the *monotonic-gene-set* included in the two filtered sets.

**Figure 4:**
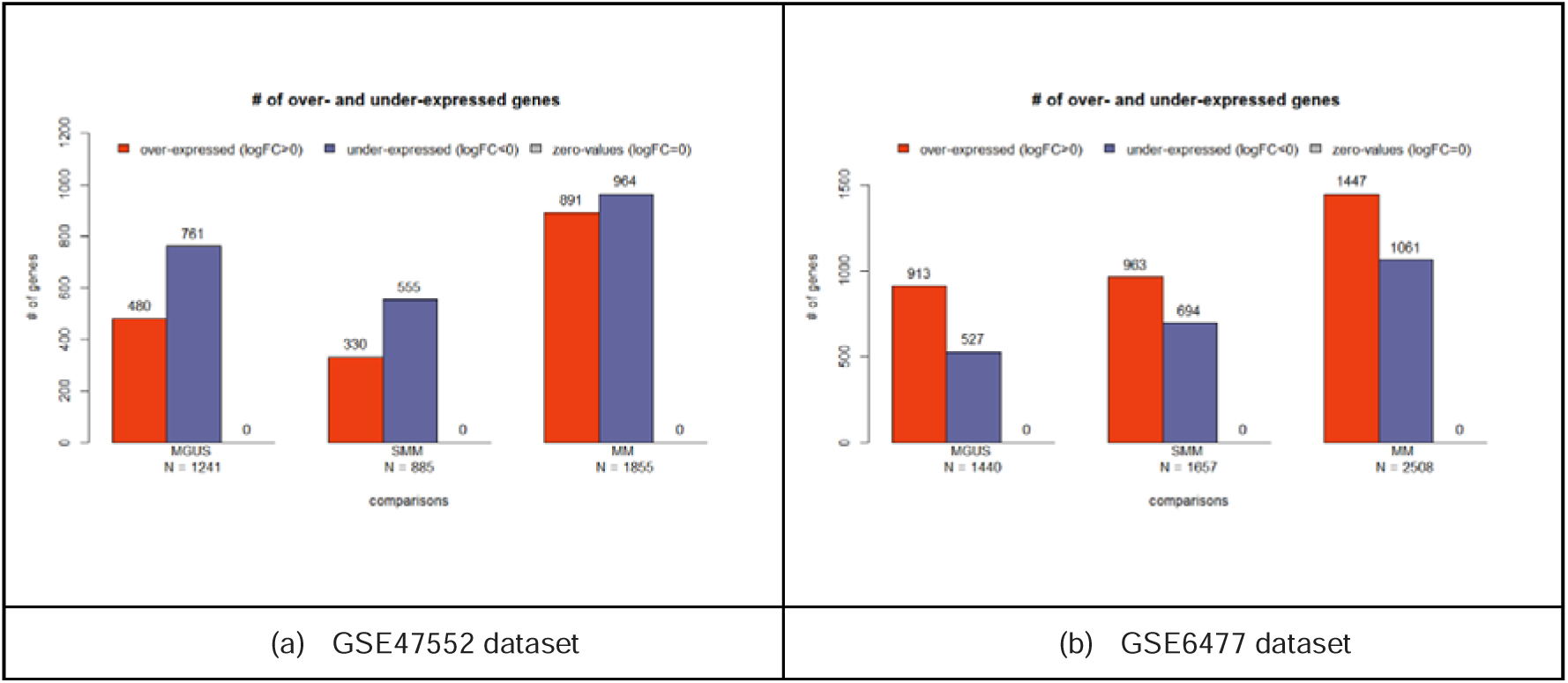
Number of over- and under-expressed genes of the monotonic-gene-set included in the two datasets under study.

### Comparison between the traditional and the monotonic differential expression analysis

In order to evaluate the statistical significance of the monotonic genes with those obtained through the traditional DEA, we performed a multi-step filtering for various thresholds th of absolute log_2_FC parameter (|log_2_FC| ?: th), where on each step we calculated the average adjusted p-value of the remaining genes. This was done in the prospect to examine whether the *monotonic-gene-set* provides higher mean-statistical significance than the *original-gene-set*. Figure 5, depicts the results of this process for each stage comparison. It is observed that the *monotonic-gene-set* exhibits slightly higher mean statistical significance than the *original-gene-set* particularly for lower thresholds of |log_2_FC| parameter. Herein we recall that the monotonic approach, although does not account for statistical significance, provides a natural parameter that shows the direction of change in the expression of a gene. However the increased mean significance at lower thresholds of |log_2_FC| parameter, suggests that genes typically excluded in traditional DEA filtering, yet exhibit a monotonic characteristic, may play a crucial role across various stages of the disease.

**Figure 5:**
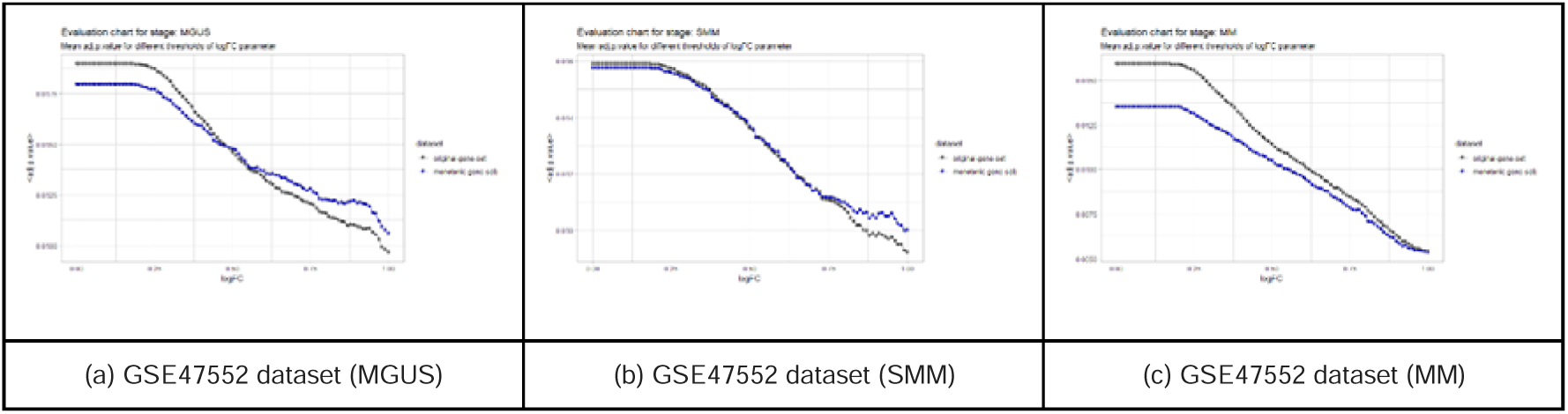
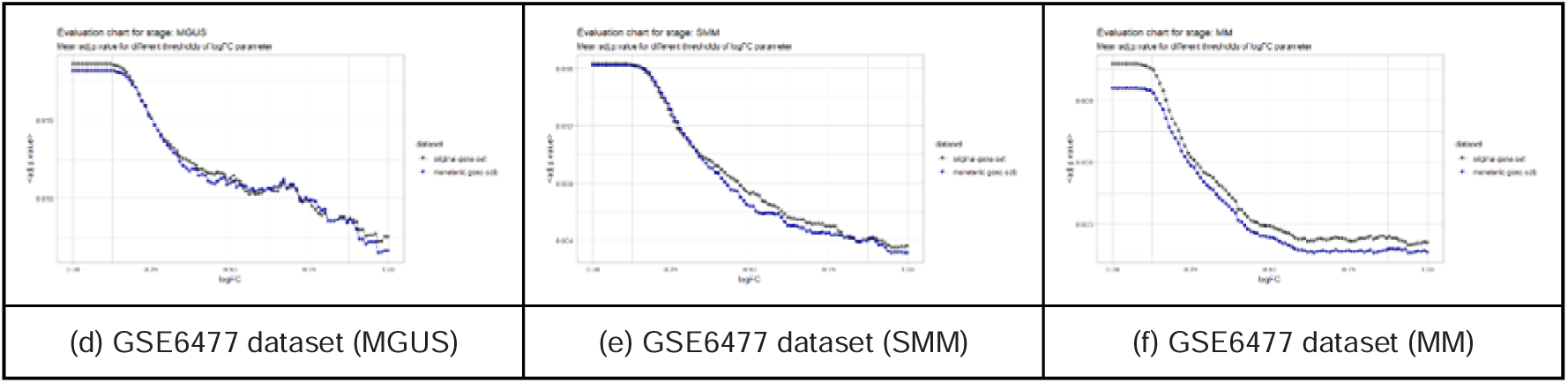
Mean adj.p.value for different thresholds of log_2_FC parameter, across different stage comparisons. (a,b,c) depict the results for the GSE47552 dataset, (d,e,f) depict the results for the GSE6477 dataset. The blue dots refer to the *monotonic-gene-set* while the grey ones refer to the *original-gene-set* accordingly.

### Developing a computational method to exploit the discrimination power of within-sample gene ratios

The preprocessing steps involved extracting the raw data without applying any normalisation or background correction algorithms. Probe IDs were further converted to EntrezIDs, to retain uniquely represented EntrezIDs, and duplicates have been merged by averaging their expression values. The common genes (n=12527) between datasets were identified, and those genes were filtered to create stage-specific gene-expression tables. The MEGs used were selected among the common genes between the datasets and across the stages after applying the adjusted p-value filtering of *p_adj_* ≤ 0.05. In consequence, for each sample we obtained the gene-expression ratios between the genes included in the sample and the identified list of MEGs accordingly, resulting in a large matrix of ratios per sample. The following equations describe this process:

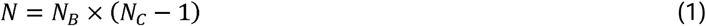

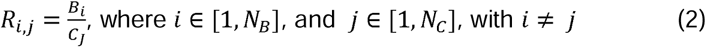

where R refers to the ratio for a pair of genes i and j, B is the expression value of each MEG included in the dataset, C refers to the expression of each gene included in the dataset, N is the total number of obtained ratios per sample, N_B_ refers to the number of identified MEGs, and N_C_ is the number of genes included in the two datasets. The obtained matrices (per sample) were merged according to their disease stage, resulting in specific lists of ratio distributions, per gene pair. In order to examine which ratio distributions can adequately separate between disease stages, we used specific statistical criteria as follows. We first examined which ratio distributions followed a normal distribution by means of the Shapiro Test [14] (ST) and kept only those with p-value > 0.05. The retained distributions were further filtered by means of their Coefficient of Variation (CV) [15] and those that showed lower than 10% dispersion around the mean were kept. From the 331599 Healthy, 117557 MGUS, 74121 SMM and 63306 MM ratios that were retained, only the 11999 were present in all four stages and kept for forward analysis.

Expanding on this approach, we used the Mann-Whitney U test [16] (MWU) (also known as the Wilcoxon rank-sum test), to examine which distributions from the remaining 11999 ratios exhibit significant statistical differences between all the possible stage comparisons. The choice of this test was driven by the nature of the data under analysis. Characteristically, the gene ratio data may not be necessarily optimally normally distributed even if we have already performed the ST, CV tests, since outliers may have a significant impact on the results. Therefore, employing a nonparametric test like the MWU allows for a more robust analysis that is less sensitive to the underlying distribution of the data. By comparing the ranks of values in the two samples rather than the values themselves, the MWU test can detect differences in the central tendency of the gene ratio distributions, making it an appropriate choice for this analysis. To this extent, using the Benjamini & Hochberg [17] approach to convert the obtained p-values, into more robust adjusted p-values, we finally performed a threshold of *p_adj_* ≤ 0.05 to finalise our filtering methodology.

Table 2 shows the number of remaining ratios (gene-pairs) after performing the above filtering process. It is observed that pairwise comparisons within disease stages seem to have much less significant pairs compared to healthy-disease stage pairwise comparisons.

Drawing from the final pairwise data shown in Table 2, two discretesets of data normally used in classification schemes were conducted: (a) the internal set of data used for training, and (b) the external set of data used for examining and evaluating the performance of the proposed classifier. Specifically, the internal set of data was built in the form of Table 1, using the pairwise ratio-distributions extracted from the two reference datasets (GSE47552 & GSE6477), which include all the required stages of Multiple Myeloma. In consequence, the external set of data was built in the same form, drawing information from the remaining five datasets included in Table 1.

**Table 1.** Summary of the transcriptomic datasets used in this study.

| Disease | GSE47552 | GSE6477 | GSE13591 | GSE14230 | GSE235356 | GSE5900 | GSE80608 |
| --- | --- | --- | --- | --- | --- | --- | --- |
| HEALTHY | 5 | 15 | 5 | 5 | 0 | 22 | 10 |
| MGUS | 20 | 22 | 11 | 5 | 319 | 44 | 10 |
| MM | 41 | 73 | 133 | 5 | 39 | 0 | 10 |
| SMM | 33 | 24 | 0 | 0 | 0 | 12 | 0 |

The underlying classification methodology reads as follows. For each ratio r obtained from a pair of genes included in the samples across the datasets under study, we firstly find the related ratio-distributions included in the internal dataset for the specific pair. Using the Maximum-Likelihood Fitting of univariate distributions we then calculate the estimated mean values (µ_A_, µ_B_) and estimated standard deviations (a_A_, a_B_) for each one of the two ratio-distributions. In consequence, we obtain two density values (d_A_, d_B_) for the given ratio r, based on normal distributions with mean equal to µ_A_, µ_B_and standard deviation equal to a_A_, a_B_, accordingly. Finally, the examined ratio is classified to that class (stage) with the maximum density. Herein each ratio is considered as a separate classifier examined across samples, where the ACC and Area Under the Curve (AUC) [18] parameters are obtained to evaluate/rank its performance. This process was performed for all the available ratios included in both the internal and external datasets, across all the available samples. The results from the analysis above are depicted in Supplementary File 3.

## RESULTS

To examine the most highly discriminative gene-pairs within the two datasets, we conducted lists of gene-pairs where: (a) wrongly classified ratios were removed, and (b) ratios with overall ACC and AUC less than 70%, 80% and 85%, have been also removed, from the internal dataset while for the external one we considered only the ACC parameter, due to the small number of significant ratios. Figure 6 depicts the distribution of the ACC and AUC parameters across gene-pairs (ratios) obtained from this classification approach, for all the six pairwise stage-comparisons. It is observed that the internal dataset exhibits much higher ACC and AUC values compared to the external dataset. In effect, this indicates that the external dataset clearly shows a weakness in contributing added value to the overall results, while the majority of the gene-pair information mainly comes from the internal dataset. However, since the goal of this classification approach is not to provide an optimal classification scheme but rather to use it as a method for identifying highly discriminative gene-pairs, this weakness is considered an acceptable trade-off for contributing added value from limited RNA-Seq experiments, since there is a quite acceptable number of gene-pairs with high accuracy ACC ?: 70%, which derive from the external dataset.

**Figure 6:**
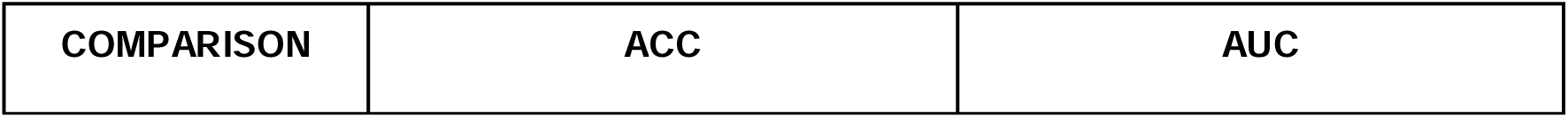
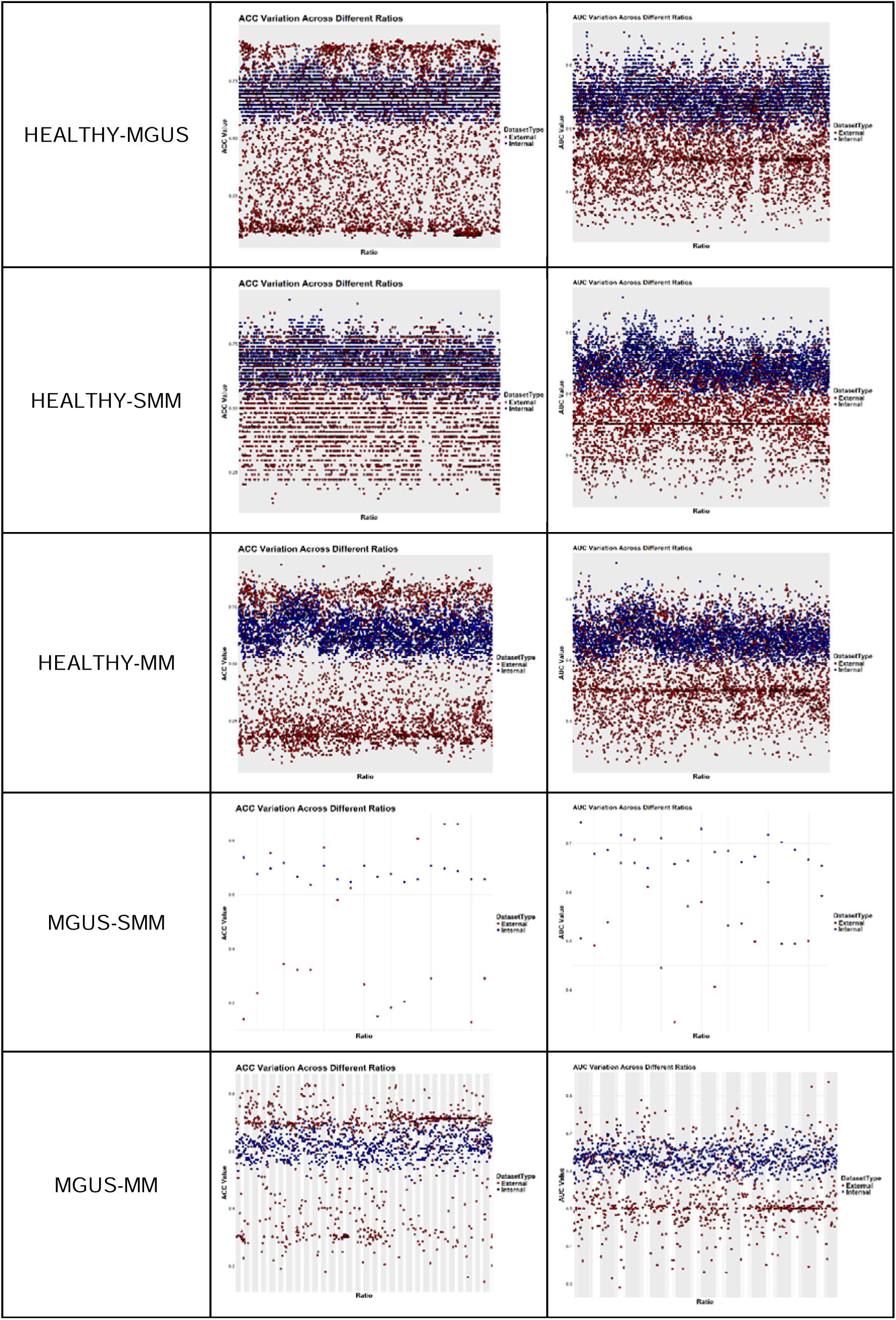
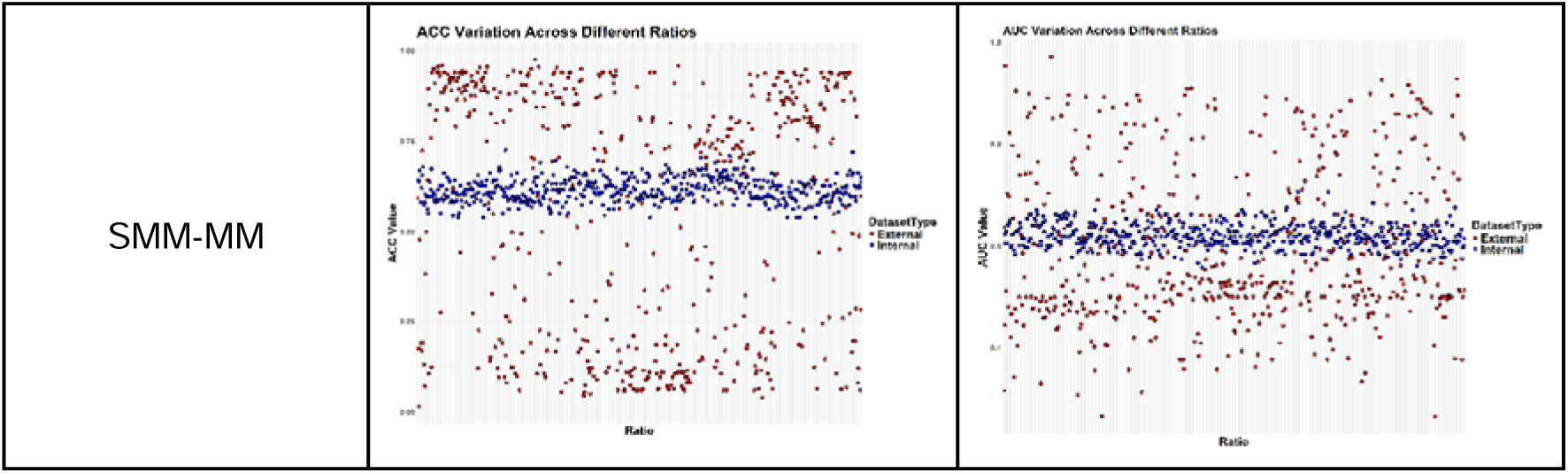
Variation of the ACC and AUC classification parameters across gene-pairs obtained from this classification approach, for all the six pairwise comparisons. The blue points refer to the internal dataset, while the red ones refer to the external dataset, accordingly.

In consequence, we further kept only the common pairs across the internal and external datasets to keep the gene-pair information that covers as much as more RNA bulk experiments. Table 3 shows the remaining gene-pairs for different thresholds of accuracy. It is observed that the performance of pairwise comparison within disease stages was poor and according to the aforementioned ratio filterings. Characteristically, only one ratio remained for within disease stage comparisons when performing the ACC ?: 70% threshold, while for larger thresholds of ACC only the healthy-vs-disease stage comparisons revealed highly discriminative gene-pairs.

The gene-lists generated for each threshold and used for the pathway-based analysis are depicted in Supplementary File 4.

Table 3 indicates that the thresholds of ACC ?: 70% and AUC ?: 70% as well as the thresholds of ACC ?: 80% and AUC ?: 80% accordingly, can adequately provide lists of gene-pairs towards examining their importance to the stages of Multiple Myeloma. On these grounds, the obtained genes were further examined at pathway level using the EnrichR R package [19], widely used for pathway enrichment analysis. Specifically, we created three lists of gene pairs as follows: (a) the *control_disease_70* which refers to genes included in the healthy vs disease gene-pairs filtered with 70% parameter thresholds, (b) the disease_disease_70 which refers to the genes included in the disease vs disease gene-pairs filtered with 70% parameter thresholds, (c) the *control_disease_80* which refers to genes included in the healthy vs disease gene-pairs filtered with 80% parameter thresholds, and the *control_disease_85* which refers to genes included in the healthy vs disease gene-pairs filtered with 85% parameter thresholds.

Figure 7 depicts the heatmap of the genes derived from the control_disease_80 list. It is observed that the majority of those genes have the same expression trend across the two datasets. In addition from the control_disease_85 list, we obtained 7 candidate genes derived from the Healthy vs MGUS stage comparison, namely the ADCY9, KRT3, GRK1, LDB3, TBXA2R, SSTR5, and the SLC44A4 gene.

**Figure 7:**
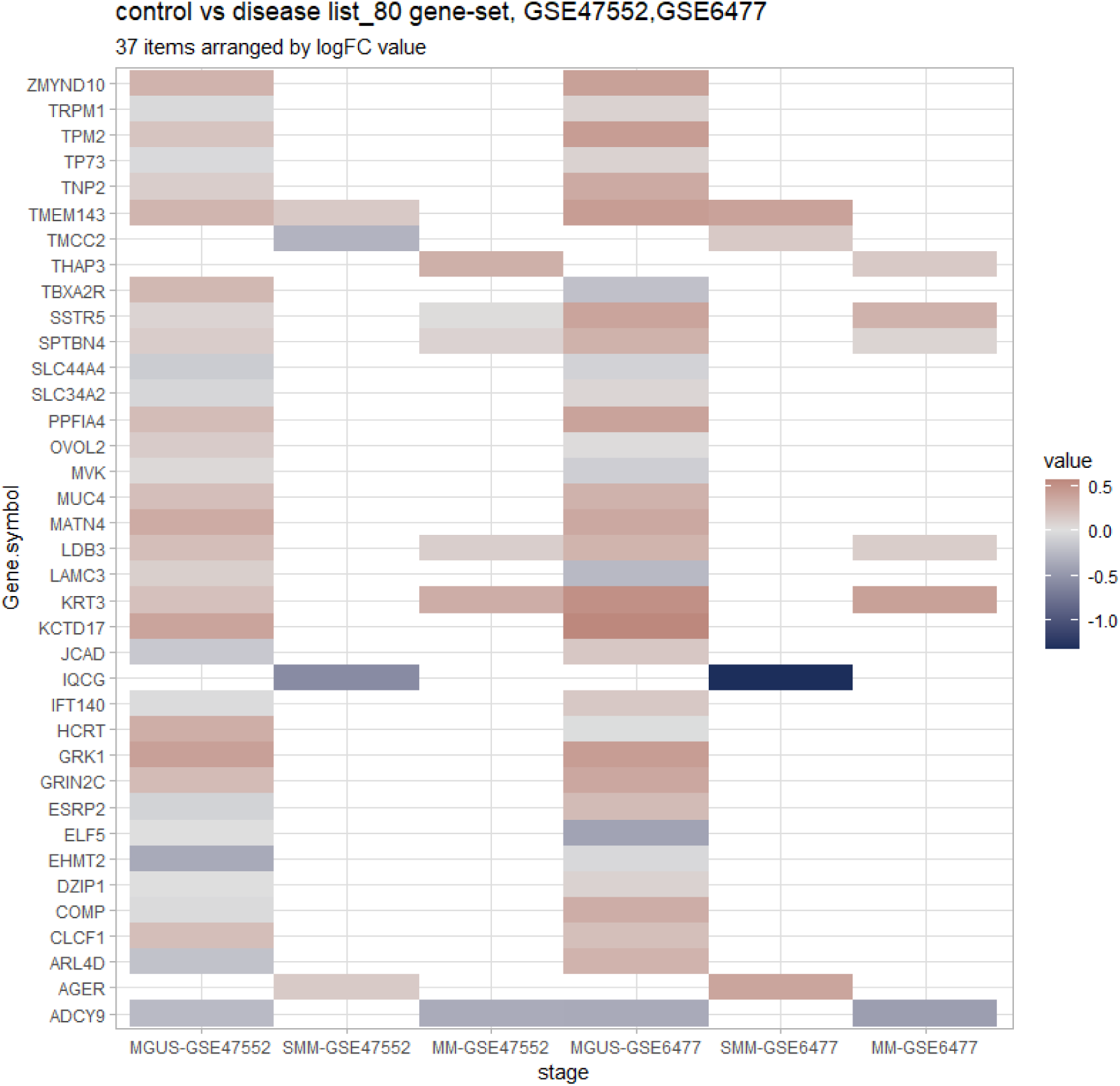
Heatmaps depicting the list of genes included in the control_disease_80 list, and their expression status across stages, in the GSE6477 and the GSE47552 datasets, accordingly. The different colours refer to the value of the log_2_FC parameter. For the rest of the lists please refer to the supplementary files.

Figure 8 depicts the top 10 pathways per stage pair, ranked by means of the *combined.score* metric provided by the EnrichR package for the “GO_Biological_Process_2023” repository. Pathways refer to the control_disease_80 list.

**Figure 8:**
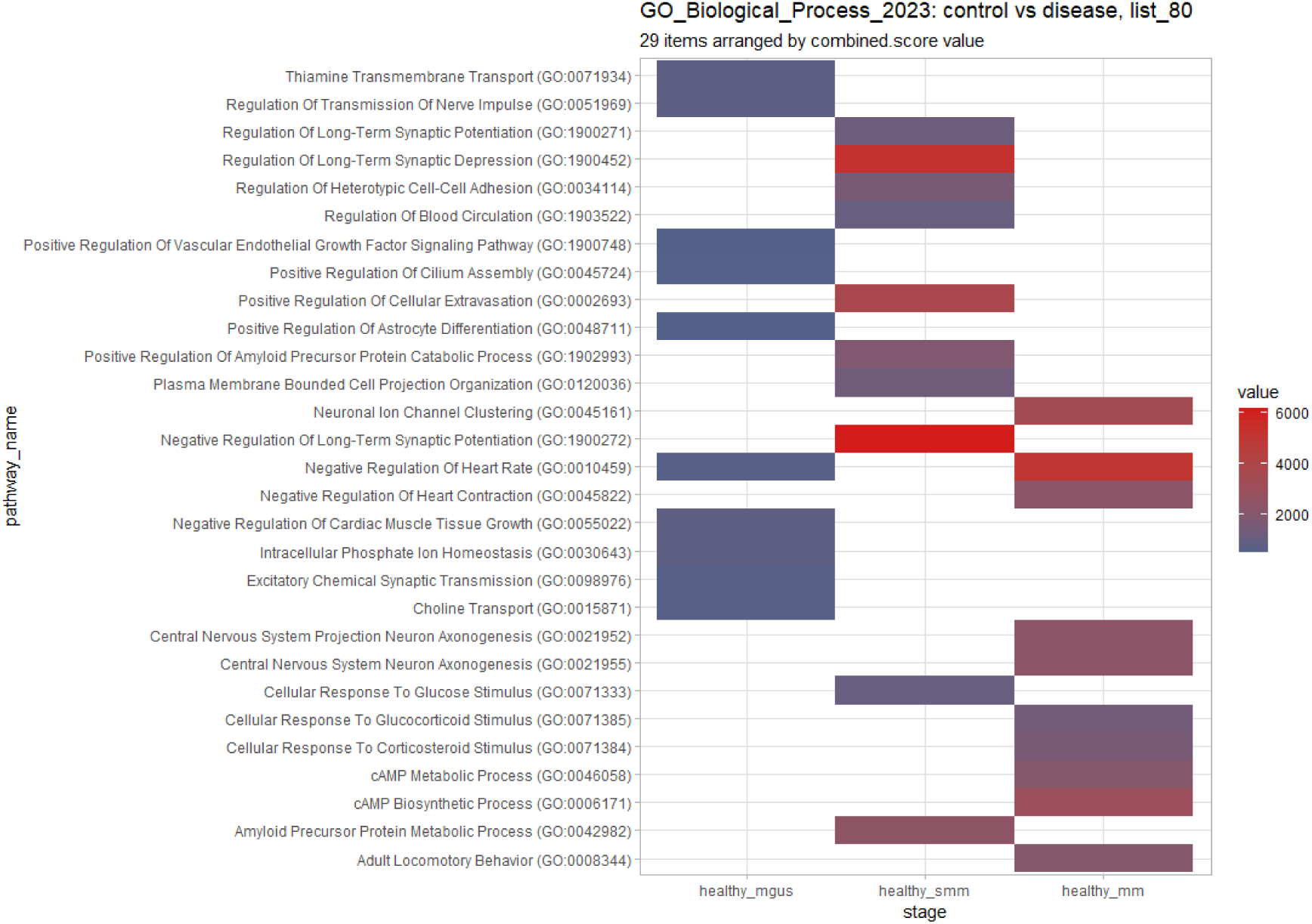
Heatmap depicting the lists of top 10 pathways per stage pair, ranked by the combined.score metric provided by the EnrichR package. The “GO_Biological_Process_2023” repository was used. The colours represent the variation of the combined.score provided by the EnrichR package. The heatmap refers to pathways obtained from the control_disease_80 list of genes described in the text.

Figure 9, depicts the top 10 pathways per stage pair, ranked by means of the *combined.score* provided by the EnrichR package, by means of the “GO_Biological_Process_2023” repository. Pathways refer to the control_disease_70 and disease_disease_70 list accordingly. It appears that the majority of pathways identified from the control_disease_80 gene list constitute a subset of those identified from the control_disease_70 gene list.

**Figure 9:**
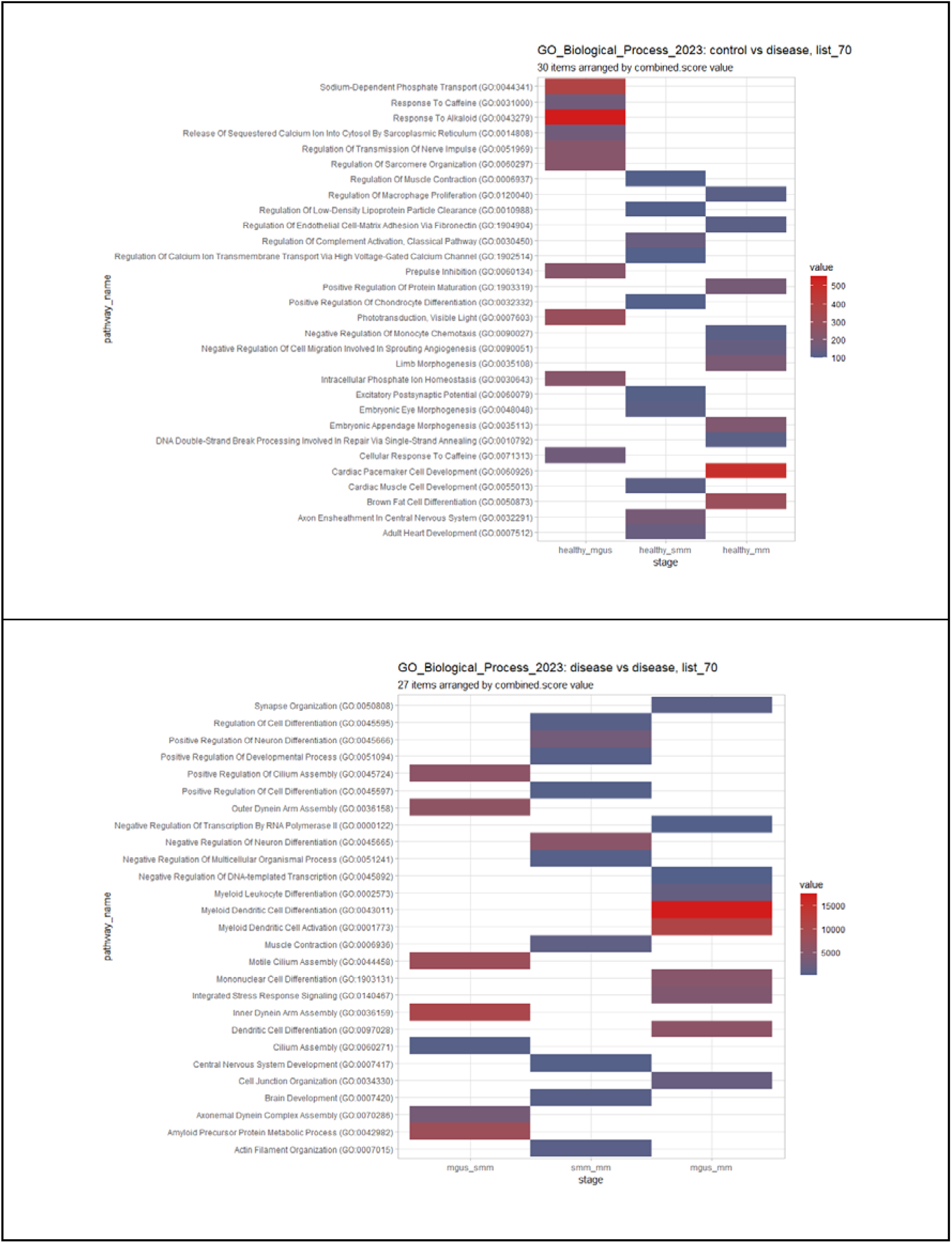
Heatmaps depicting the lists of top 10 pathways per stage pair, ranked by means of the combined.score provided by the EnrichR package. The “GO_Biological_Process_2023” repository was used. The colours represent the variation of the combined.score provided by the EnrichR package. The top heatmap refers to pathways obtained from the control_disease_70 list of genes described in the text. The bottom heatmap refers to pathways obtained from the disease_disease_70 list of genes described in the text.

## Discussion

Multiple Myeloma is the second most common haematological malignancy and is estimated that 35,780 new cases will be created in 2024, which is equal to 1.8% of all new cancer cases based on the National Cancer Institute (NIH). According to the same institute, the 5-year relative survival was equal to 61.1% according to data during the period 2014-2020 [20]. To understand the complexity and molecular profiling of Multiple Myeloma (MM), this study provides a comprehensive analysis of gene expression changes and pathway disruptions across MM and its precursor states, MGUS and SMM. Based on the methodology introduced by Bourdakou, et al. (2021), our analysis identified distinct gene expression profiles for MGUS, SMM, and MM stages in two datasets, namely the GSE6477 and the GSE47552.

The subsequent reduction in DEGs after performing the adjusted p-value and log_2_FC filtering thresholds, underscored the dynamic nature of gene expression across disease stages. Genes previously associated with MM were identified through differential expression analysis across the two datasets. Specifically, Cyclin D1 (CCND1) [1], has been strongly associated with MM due to the IgH Translocation t(11;14) (q13;q32), which was found up-regulated in MGUS. Cyclin D2 (CCND2) together with CCND1 takes part in the cell-cycle mechanism affecting cell-cycle arrest, DNA repair and apoptosis, and was also found upregulated in MGUS. In SMM, Baculoviral IAP Repeat Containing 2 (BIRC2), a gene associated with the NF-K pathway as well as KRAS, which takes part in the MAPK signalling pathway [21], was found up-regulated in our study. Baculoviral IAP Repeat Containing 3 (BIRC3) and NFKB Inhibitor Alpha (NFKBIA), genes associated with the NF-K pathway, were found to be dysregulated in both SMM and MM. MAF BZIP Transcription Factor B (MAFB) which is mentioned as a primary abnormality for MM due to the IgH Translocation t(14;20) (q32;q11) was found downregulated in both stages. DNA Damage-related genes ATM Serine/Threonine Kinase (ATM) and Tumor Protein P53 (TP53) were down-regulated in MM. Deletion of 17p (Del(17p)), previously associated with MM, was linked with the dysregulation of TP53. In both MGUS, SMM and MM stages, the expressions of RB Transcriptional Corepressor 1 (RB1) and CD27 Molecule (CD27), were downregulated. RB1 is part of the cell-cycle mechanism and is associated with Del(13q), and CD27, was previously associated with Del(12p) [1,21,22].

Genes with characteristic monotonic trends were then found to increase our understanding of the factors causing the progression from the asymptomatic and premalignant stages to active Multiple Myeloma. To our understanding, this was the first time a gene monotonic signature was found for MM with the monotonicity analysis to be able to further elucidate the dynamics shown in DEGs, revealing genes with consistent expression trends, which could serve as potential indicators for disease progression. Three of those RB1, CD27 and TP53 were mentioned above for their association as DEGs while the MCL1 Apoptosis Regulator, BCL2 Family Member (MCL1) gene, previously associated with Multiple Myeloma because of 1q Gain [23], was found in our analysis through monotonicity analysis.

To refine our understanding of how those MEGs can be used to indicate the disease stages, we calculated MEG-associated ratios, identifying those characterised by normal distribution and small dispersion. The analysis demonstrated that several ratios effectively distinguished healthy from disease samples, though this distinction was less clear in disease-to-disease comparisons due to overlapping distributions. In consequence, we proceeded with performing pairwise comparisons. Mann-Whitney U test was used to calculate the difference between the two distributions and based on the *p_adj_* < 0.05 threshold a final list of ratios for each comparison was created (Table 2). To those lists, we can see a decreased trend in ratios as the disease progresses, from 4402 (Healthy-MGUS) to 4089 (Healthy-SMM) and 3621 (Healthy-MM) showing the complexity of this disease as it progresses. This is also supported by the within disease stage comparisons where only 19 ratios were significant for MGUS-SMM comparison while it increases to 535 for SMM-MM and 822 for MGUS-MM. Based on the above we can see that the ratio profiles of MGUS and SMM are closer and as the disease progresses the ratio profiles move closer to MM. The remaining gene pair ratios were then measured to an internal and external dataset in an attempt to use gene pair ratios with specific characteristics to distinguish samples between different disease conditions. Our results highlighted the biology behind the progression of Multiple Myeloma, where according to the aforementioned filterings, the gene pair ratios between control-to-disease comparisons were more informative than those from within disease stage comparisons due to the complexity and similarity of their molecular profiles. However, we have to note that, the small sample size of only two datasets used to find possible gene pair ratios and the unbalanced sample size of the external dataset (only 12 SMM samples) may have interfered with our results.

**Table 2:** Number of remaining gene-pairs for all possible stage comparisons, after performing the filtering methodology described in the text.

| Stage A | Stage B | # gene-pairs (ratios) |
| --- | --- | --- |
| Healthy | MGUS | 4402 |
| Healthy | SMM | 4089 |
| Healthy | MM | 3621 |
| MGUS | SMM | 19 |
| MGUS | MM | 822 |
| SMM | MM | 535 |

**Table 3:** Final number of ratios for each pairwise comparison

| Stage A | Stage B | # gene-pairs (ratios) | # gene-pairs (ratios) | # gene-pairs (ratios) |
| --- | --- | --- | --- | --- |
| **INTERNAL FILTERING** | | $ACC \geq 70\%$ | $ACC \geq 80\%$ | $ACC \geq 85\%$ |
| | | $AUC \geq 70\%$ | $AUC \geq 80\%$ | $AUC \geq 85\%$ |
| **EXTERNAL FILTERING** | | $ACC \geq 70\%$ | $ACC \geq 80\%$ | $ACC \geq 85\%$ |
| **Healthy** | **MGUS** | **410** | **23** | **4** |
| **Healthy** | **SMM** | **274** | **2** | **0** |
| **Healthy** | **MM** | **148** | **4** | **0** |
| **MGUS** | **SMM** | **1** | **0** | **0** |
| **MGUS** | **MM** | **1** | 0 | 0 |
| **SMM** | **MM** | **1** | 0 | 0 |

Enrichment analysis performed on the final lists of gene-pairs, highlighted critical biological processes and pathway disruption across Multiple Myeloma stages, as shown in Figures 8,9. Characteristically, Calcium-related pathways related to bone disease, that have been previously associated with Multiple Myeloma [24], were enriched in MGUS and SMM using list_70 gene-pairs. Indeed, Li et. al, (2021), mentioned that 70-80% of MM patients suffer from Myeloma bone disease, a condition accompanied with fractures and hypercalcemia. The authors also state that patients with irreversible kidney damage and increased renal tubular calcium reabsorption show elevated serum calcium and abnormal bone remodelling [25]. Furthermore, glucocorticoids (GCs) were highlighted in MM list_80. According to Bayraktar (2010), GCs may rapidly reduce paraprotein production, reduce inflammation and fibrosis in renal parenchyma, and decrease serum calcium levels in newly diagnosed MM patients [26]. GCs regulate many physiological functions such as stress responses, metabolism (lipid homeostasis), immunosuppressive effects (impacting dendritic cells and proinflammatory T lymphocytes, apoptosis) and hepatic functions by maintaining glucose homeostasis (gluconeogenesis and glycogenolysis), as well as growth, development and cell differentiation [27]. Cardiac and muscle involvement were also highlighted in both list_70 and list_80. Proskuriakova et.al (2021) stated that Multiple Myeloma is associated with light-chain amyloidosis (AL) that causes both toxic and infiltrative effects on the heart. This effect is associated with the appearance of amyloid cardiomyopathy, which is characterised by the accumulation of fibrils in atria, ventricles and vessels causing impaired cell metabolic processes. Also due to toxicity effects that occur, it can lead to heart failure through autophagy, accumulation of reactive oxygen species (ROS) and disturbance of mitochondria and lysosomes [28]. Another system that was found to be impaired according to our enrichment analysis is the neuron system. Peripheral neuropathy (PN) has already been established for its association with monoclonal gammopathies due to cell plasma dyscrasia [29]. Even though PN can be induced in patients after treatment studies have already found that it can appear in newly diagnosed MM patients. Several mechanisms can result in the appearance of PN such as the M protein accumulation, deposits of light-chain immunoglobulins in nerves and events caused by myeloma bone disease that could directly damage nerve cells. Also, hyperviscosity of blood due to the high levels of M-protein could cause neurological symptoms due to the reduced flow of blood [30].

We recall that Figures 8 and 9 show that the majority of pathways identified from the control_disease_80 gene list constitute a subset of those identified from the control_disease_70 gene list, which in turn contains much more highly ranked genes. Therefore, we estimate that the most highly filtered candidate genes for stage evaluation are likely to be those included in the control_disease_80 and control_disease_85 gene list. A brief review of the literature on these genes revealed that most of them have been previously linked to various types of cancer, with a significant number showing both direct and indirect associations with Multiple Myeloma. In effect, this finding reinforces the reliability of the proposed methodology. In the following, we present a summary of the literature review further providing arguments that relate the underlying genes with the stages of Multiple Myeloma.

Starting with the genes included in the control_disease_85 list, the ADCY9 gene has been found in the list of candidate genes related to hereditary neuropathies. Anderson et al. (2019) mentioned that ADCY9 was one of 11 differentially expressed proteins identified within the t(11;14) subgroup of Human Myeloma Cell Lines (HMCLs) [31]. Additionally, Jérôme Moreaux et al. (2011) using HMCLs with MMSET/FGFR3 translocations, were able to identify through DEA the overexpression of ADCY9 [32]. Lastly, ADCY9 has been associated also with other types of cancers: (a) overexpressed in colon cancer [33], (b) acts as a tumour suppressor to inhibit proliferation, migration, and invasion in lung adenocarcinoma [34], (c) associated with the risk for hepatocellular carcinoma [35].

KRT3 has been mentioned among genes aberrantly methylated in myeloma [36]. Li et al. (2022) identified KRT3 expression in high- and low-risk patients with bladder cancer based on a large population cohort while Liang et al (2022) mentioned that KRT3 could serve as a potential biomarker for monitoring oral leukoplakia (OLK) and promote the early diagnosis of oral squamous cell carcinoma (OSCC) [37,38]. GRK1 is part of the GRK family, which consists of seven members (GRK1-7). Among them, GRK6 has been found to be associated with Multiple Myeloma [9].

LIM Domain Binding 3 (LDB3) is a striated muscle-specific Z-band protein involved in mechanosensory actin cytoskeleton remodelling [39]. LDB3 was previously selected to establish an overall survival (OS) prediction model in soft tissue sarcoma and was also included in an 11-gene prognostic signature for uveal melanoma metastasis [40,41]. Wang et al. (2021) after applying Quercetin in Multiple Myeloma cells, identified that the expression of Thromboxane A2 Receptor (TBXA2R) was reduced, highlighting the potential role of this receptor in Multiple Myeloma cells [42].

Somatostatin Receptor 5 (SSTR5) isn’t a well-established gene in cancer research, but a 2003 study demonstrated that SSTR5 was expressed in human uveal melanomas, with immunohistochemical staining showing positivity in 58% of the tumours [43]. Qin et al (2022), created a ferroptosis gene signature using machine-learning approaches and identified that the SLC44A4 gene appears to correlate with the signature after applying it to MM samples [44].

Moving to the control_disease_80 lists, several directly or indirectly related genes to Multiple Myeloma were uncovered. Starting with, KCTD17 somatic mutations were related to high-risk acute Myeloid Leukaemia characterised by -7/del(7q) [45]. SPTBN4 has been correlated with axonal neuropathy causing myopathy and hypoacusis [46] while THAP3, has been correlated with the CD45 expression that promotes apoptosis in chemotherapy patients with Myeloid Leukaemia [47]. Furthermore, TMCC2 has been mentioned by Ping Zhou et al. (2023) who performed single-cell RNA sequencing (scRNA-seq) employing the 10X Genomics Chromium Single Cell 3′ method on Bone Marrow Interstitial Light-Chain Amyloid samples in patients with myeloma. The authors showed significantly up-regulated erythroid genes involved in oxygen transport, including TMCC2 [48].

Tumor protein p73 (TP73) methylations, deletions and point mutations are common in MM, decreasing the capability of the organism to activate apoptosis or senescence of DNA damage cells [49]. Gillardin et al. (2017) tried to re-activate TP73 using Decitabine and Melphalan in TP53 Deficient Myeloma Cells and bypass the inhibition of TP53 but the combination of those compounds failed [50]. Furthermore, a study in 2010, after conducting DNA methylation analysis in bone marrow DNA samples of 21 MGUS 44 MM patients revealed that 33% and 45% respectively, had TP73 methylations suggesting this as an early event in the pathogenesis and development of Plasma Cell disorders [51].

Euchromatic histone-lysine N-methyltransferase 2 (EHMT2) copy number amplifications that encode the histone methyltransferase G9a, have been frequently detected in MM [52,53]. Interesting results by Ishiguro et al. (2021) depict that dual inhibition of EHMT2/G9a reduced H3K27/H3K9 methylation levels in MM cells and increased expression of endogenous retrovirus (ERV) genes, suggesting that activation of ERV genes may induce the IFN response [54].

Despite the limited number of gene pairs in the disease_disease_70 list, these specific genes have already been associated with Multiple Myeloma. For instance, the TPM2 has been mentioned by Dunphy et al. (2023) who performed survival analysis on gene expressions from patients groups with MM. The authors found that there is a trend towards decreased overall survival in those with the high expression of six (TAGLN2, CA2, ITGA2, LGALS1, TPM2, TMOD3) out of the seven proteins analysed [55]. Then, the expression of DLX2, a differentiation-regulating gene, was inhibited because of the upregulation of RUNX2 which promotes the suppression of myeloma osteoblast activity [56]. Lastly, Guang Yang et al.(2018), who performed wholeLexome sequencing on 44 multiple myeloma patients, identified the PPFIA4 as the nearest gene, among those listed in the top SNPs obtained from the Exome-Wide Association Analysis [57].

## Conclusion

This study elucidates the gene expression dynamics and pathway disruptions in three generic stages of Multiple Myeloma. Traditional differential expression analysis identified several significant genes including the *CCND1*, *CCND2*, *BIRC2*, *BIRC3*, *KRAS*, *MAFB*, *ATM*, *TP53*, *RB1*, and *CD27*. Utilising a recently introduced computational methodology designed to detect MEGs, we further identified the *RB1*, *CD27*, *TP53*, and *MCL1* monotonic genes exhibiting a consistent expression trend across stages. Evaluation and testing, revealed that applying the traditional differential analysis filtering on monotonic gene-set, yielded slightly higher statistically significant genes and pathways related to Multiple Myeloma. On these grounds, we further proposed a ranking methodology based on monotonic gene pairs across samples, which effectively identified 37 candidate genes, potentially associated with specific stages of Multiple Myeloma, demonstrating high accuracy in distinguishing between control and disease samples.

Enrichment analysis by means of those genes uncovered significant groups of pathways linked to generalised mechanisms that could potentially be affected at each stage of Multiple Myeloma. Even more, literature exploration performed on the identified genes, revealed that most of them have been previously linked to various types of cancer, with a significant number showing both direct and indirect associations with Multiple Myeloma. Future steps include applying the underlying methodology to other diseases in the prospect of examining its potential in recognising gene-pairs that may not be highlighted by classical analysis but are critical for disease progression.

## Author Contributions

Methodology, G.G., G.M., N.K., M.M.B., E.A and G.M.S; validation, G.G. and K.S.; investigation, G.G. and G.M., writing—original draft preparation, G.G.; writing—review and editing, G.G., G.M., K.S., N.K., M.M.B., E.A and G.M.S.; visualization, G.G. and G.M..; supervision, G.M. and G.M.S.; project administration, G.M.S.

## Glossary

GEO: Gene Expression Omnibus
DEA: Differential Expression Analysis
DEGs: Differential Expressed Genes
MEGs: Monotonically Expressed Genes
MEPs: Monotonically Enriched Pathways
MGUS: Monoclonal Gammopathy of Undetermined Significance
SMM: Smoldering Multiple Myeloma
MM: Multiple Myeloma
ACC: Accuracy
AUC: Area Under Curve

## References

[1] Castaneda O, Baz R. Multiple Myeloma Genomics - A Concise Review. Acta Medica Acad 2019;48:57–67. 10.5644/ama2006-124.242.

[2] Bourdakou MM, Spyrou GM, Kolios G. Colon Cancer Progression Is Reflected to Monotonic Differentiation in Gene Expression and Pathway Deregulation Facilitating Stage-specific Drug Repurposing. Cancer Genomics Proteomics 2021;18:757–69. 10.21873/cgp.20295.

[3] Edgar R, Domrachev M, Lash AE. Gene Expression Omnibus: NCBI gene expression and hybridization array data repository. Nucleic Acids Res 2002;30:207–10. 10.1093/nar/30.1.207.

[4] Davis S, Meltzer PS. GEOquery: a bridge between the Gene Expression Omnibus (GEO) and BioConductor. Bioinforma Oxf Engl 2007;23:1846–7. 10.1093/bioinformatics/btm254.

[5] Agnelli L, Mosca L, Fabris S, Lionetti M, Andronache A, Kwee I, et al. A SNP microarray and FISH-based procedure to detect allelic imbalances in multiple myeloma: an integrated genomics approach reveals a wide gene dosage effect. Genes Chromosomes Cancer 2009;48:603–14. 10.1002/gcc.20668.

[6] Ria R, Todoerti K, Berardi S, Coluccia AML, De Luisi A, Mattioli M, et al. Gene expression profiling of bone marrow endothelial cells in patients with multiple myeloma. Clin Cancer Res Off J Am Assoc Cancer Res 2009;15:5369–78. 10.1158/1078-0432.CCR-09-0040.

[7] Sun F, Cheng Y, Ying J, Mery D, Al Hadidi S, Wanchai V, et al. A gene signature can predict risk of MGUS progressing to multiple myeloma. J Hematol OncolJ Hematol Oncol 2023;16:70. 10.1186/s13045-023-01472-y.

[8] López-Corral L, Corchete LA, Sarasquete ME, Mateos MV, García-Sanz R, Fermiñán E, et al. Transcriptome analysis reveals molecular profiles associated with evolving steps of monoclonal gammopathies. Haematologica 2014;99:1365–72. 10.3324/haematol.2013.087809.

[9] Zhan F, Barlogie B, Arzoumanian V, Huang Y, Williams DR, Hollmig K, et al. Gene-expression signature of benign monoclonal gammopathy evident in multiple myeloma is linked to good prognosis. Blood 2007;109:1692–700. 10.1182/blood-2006-07-037077.

[10] Chng WJ, Kumar S, Vanwier S, Ahmann G, Price-Troska T, Henderson K, et al. Molecular dissection of hyperdiploid multiple myeloma by gene expression profiling. Cancer Res 2007;67:2982–9. 10.1158/0008-5472.CAN-06-4046.

[11] McNee G, Eales KL, Wei W, Williams DS, Barkhuizen A, Bartlett DB, et al. Citrullination of histone H3 drives IL-6 production by bone marrow mesenchymal stem cells in MGUS and multiple myeloma. Leukemia 2017;31:373–81. 10.1038/leu.2016.187.

[12] Ritchie ME, Phipson B, Wu D, Hu Y, Law CW, Shi W, et al. limma powers differential expression analyses for RNA-sequencing and microarray studies. Nucleic Acids Res 2015;43:e47. 10.1093/nar/gkv007.

[13] Gordon K Smyth. Linear models and empirical bayes methods for assessing differential expression in microarray experiments. Stat Appl Genet Mol Biol 2004;3. 10.2202/1544-6115.1027.

[14] Shapiro SS, Wilk MB. An analysis of variance test for normality (complete samples). Biometrika 1965;52:591–611. 10.1093/biomet/52.3-4.591.

[15] Kesteven GL. The Coefficient of Variation. Nature 1946;158:520–1. 10.1038/158520c0.

[16] Mann–Whitney Test. Concise Encycl. Stat., New York, NY: Springer New York; 2008, p. 327–9. 10.1007/978-0-387-32833-1_243.

[17] Benjamini Y, Hochberg Y. Controlling the False Discovery Rate: A Practical and Powerful Approach to Multiple Testing. J R Stat Soc Ser B Stat Methodol 1995;57:289–300. 10.1111/j.2517-6161.1995.tb02031.x.

[18] Hand DJ, Till RJ. A Simple Generalisation of the Area Under the ROC Curve for Multiple Class Classification Problems. Mach Learn 2001;45:171–86. 10.1023/A:1010920819831.

[19] Wajid Jawaid. enrichR: Provides an R Interface to “Enrichr” 2023.

[20] National Cancer Institute. Cancer Stat Facts: Myeloma. Natl Cancer Inst 2024. https://seer.cancer.gov/statfacts/html/mulmy.html (accessed March 9, 2024).

[21] Cardona-Benavides IJ, de Ramón C, Gutiérrez NC. Genetic Abnormalities in Multiple Myeloma: Prognostic and Therapeutic Implications. Cells 2021;10:336. 10.3390/cells10020336.

[22] Manier S, Salem KZ, Park J, Landau DA, Getz G, Ghobrial IM. Genomic complexity of multiple myeloma and its clinical implications. Nat Rev Clin Oncol 2017;14:100–13. 10.1038/nrclinonc.2016.122.

[23] Clarke SE, Fuller KA, Erber WN. Chromosomal defects in multiple myeloma. Blood Rev 2024;64:101168. 10.1016/j.blre.2024.101168.

[24] Oyajobi BO. Multiple myeloma/hypercalcemia. Arthritis Res Ther 2007;9 Suppl 1:S4. 10.1186/ar2168.

[25] Li T, Chen J, Zeng Z. Pathophysiological role of calcium channels and transporters in the multiple myeloma. Cell Commun Signal CCS 2021;19:99. 10.1186/s12964-021-00781-4.

[26] Bayraktar UD, Warsch S, Pereira D. High-dose glucocorticoids improve renal failure reversibility in patients with newly diagnosed multiple myeloma. Am J Hematol 2011;86:224–7. 10.1002/ajh.21922.

[27] Vitellius G, Trabado S, Bouligand J, Delemer B, Lombès M. Pathophysiology of Glucocorticoid Signaling. Ann Endocrinol 2018;79:98–106. 10.1016/j.ando.2018.03.001.

[28] Proskuriakova E, Jada K, Kakieu Djossi S, Khedr A, Neupane B, Mostafa JA. Mechanisms and Potential Treatment Options of Heart Failure in Patients With Multiple Myeloma. Cureus 2021;13:e15943. 10.7759/cureus.15943.

[29] Ballegaard M, Nelson LM, Gimsing P. Comparing neuropathy in multiple myeloma and AL amyloidosis. J Peripher Nerv Syst JPNS 2021;26:75–82. 10.1111/jns.12428.

[30] Oortgiesen BE, Kroes JA, Scholtens P, Hoogland J, Dannenberg-de Keijzer P, Siemes C, et al. High prevalence of peripheral neuropathy in multiple myeloma patients and the impact of vitamin D levels, a cross-sectional study. Support Care Cancer Off J Multinatl Assoc Support Care Cancer 2022;30:271–8. 10.1007/s00520-021-06414-3.

[31] Anderson G. Plasma membrane profiling of multiple myeloma and the identification of novel monoclonal antibody targets. Apollo - University of Cambridge Repository, 2019. 10.17863/CAM.40588.

[32] Moreaux J, Klein B, Bataille R, Descamps G, Maïga S, Hose D, et al. A high-risk signature for patients with multiple myeloma established from the molecular classification of human myeloma cell lines. Haematologica 2011;96:574–82. 10.3324/haematol.2010.033456.

[33] Li H, Liu Y, Liu J, Sun Y, Wu J, Xiong Z, et al. Assessment of ADCY9 polymorphisms and colorectal cancer risk in the Chinese Han population. J Gene Med 2021;23:e3298. 10.1002/jgm.3298.

[34] Tang Y, Wang T, Zhang A, Zhu J, Zhou T, Zhou Y-L, et al. ADCY9 functions as a novel cancer suppressor gene in lung adenocarcinoma. J Thorac Dis 2023;15:1018–35. 10.21037/jtd-22-1027.

[35] Chao X, Jia Y, Feng X, Wang G, Wang X, Shi H, et al. A Case-Control Study of ADCY9 Gene Polymorphisms and the Risk of Hepatocellular Carcinoma in the Chinese Han Population. Front Oncol 2020;10:1450. 10.3389/fonc.2020.01450.

[36] Heuck CJ, Mehta J, Bhagat T, Gundabolu K, Yu Y, Khan S, et al. Myeloma is characterized by stage-specific alterations in DNA methylation that occur early during myelomagenesis. J Immunol Baltim Md 1950 2013;190:2966–75. 10.4049/jimmunol.1202493.

[37] Guzman De Avila J, Silvera-Redondo C, Alviz-Amador A. Bioinformatic Analysis of Plus Gene Expression Related to Progression from Leukoplakia to Oral Squamous Cell Carcinoma. Asian Pac J Cancer Prev APJCP 2022;23:3833–42. 10.31557/APJCP.2022.23.11.3833.

[38] Li C, Shi Y, Zuo L, Xin M, Guo X, Sun J, et al. Identification of Biomarkers Associated with Cancerous Change in Oral Leukoplakia Based on Integrated Transcriptome Analysis. J Oncol 2022;2022:4599305. 10.1155/2022/4599305.

[39] Blech-Hermoni Y, Subedi K, Silver M, Jensen L, Coscia S, Kates MM, et al. Expression of LIM domain-binding 3 (LDB3), a striated muscle Z-band alternatively spliced PDZ-motif protein in the nervous system. Sci Rep 2023;13:270. 10.1038/s41598-023-27531-5.

[40] Li J, Dong J, Li M, Zhu H, Xin P. Potential mechanisms for predicting comorbidity between multiple myeloma and femoral head necrosis based on multiple bioinformatics 2023. 10.21203/rs.3.rs-3792368/v1.

[41] Li Y, Yang X, Yang J, Wang H, Wei W. An 11-gene-based prognostic signature for uveal melanoma metastasis based on gene expression and DNA methylation profile. J Cell Biochem 2019;120:8630–9. 10.1002/jcb.28151.

[42] Wang H, Yu D, Zhang H, Ma R, Wu H, Zhai H, et al. Quercetin inhibits the proliferation of multiple myeloma cells by upregulating PTPRR expression. Acta Biochim Biophys Sin 2021;53:1505–15. 10.1093/abbs/gmab128.

[43] Ardjomand N, Ardjomand N, Schaffler G, Radner H, El-Shabrawi Y. Expression of somatostatin receptors in uveal melanomas. Invest Ophthalmol Vis Sci 2003;44:980–7. 10.1167/iovs.02-0481.

[44] Qin J, Sharma A, Wang Y, Tobar-Tosse F, Dakal TC, Liu H, et al. Systematic discrimination of the repetitive genome in proximity of ferroptosis genes and a novel prognostic signature correlating with the oncogenic lncRNA CRNDE in multiple myeloma. Front Oncol 2022;12:1026153. 10.3389/fonc.2022.1026153.

[45] McNerney ME, Brown CD, Peterson AL, Banerjee M, Larson RA, Anastasi J, et al. The spectrum of somatic mutations in high-risk acute myeloid leukaemia with -7/del(7q). Br J Haematol 2014;166:550–6. 10.1111/bjh.12964.

[46] Finsterer J, Löscher WN, Wanschitz J, Iglseder S. Orphan Peripheral Neuropathies. J Neuromuscul Dis 2021;8:1–23. 10.3233/JND-200518.

[47] Al Barashdi MAS, Ali A, McMullin MF, Mills K. CD45 inhibition in myeloid leukaemia cells sensitizes cellular responsiveness to chemotherapy. Ann Hematol 2024;103:73–88. 10.1007/s00277-023-05520-y.

[48] Zhou P, Karagiannis T, Tai A, Ma X, Toskic D, Fogaren T, et al. Bone Marrow Interstitial Light-Chain Amyloid and Its Microenvironment. Blood 2023;142:3317–3317. 10.1182/blood-2023-178029.

[49] Petrilla C, Galloway J, Kudalkar R, Ismael A, Cottini F. Understanding DNA Damage Response and DNA Repair in Multiple Myeloma. Cancers 2023;15:4155. 10.3390/cancers15164155.

[50] Gillardin P-S, Descamps G, Maiga S, Tessoulin B, Djamai H, Lucani B, et al. Decitabine and Melphalan Fail to Reactivate p73 in p53 Deficient Myeloma Cells. Int J Mol Sci 2017;19:40. 10.3390/ijms19010040.

[51] Stanganelli C, Arbelbide J, Fantl DB, Corrado C, Slavutsky I. DNA methylation analysis of tumor suppressor genes in monoclonal gammopathy of undetermined significance. Ann Hematol 2010;89:191–9. 10.1007/s00277-009-0818-3.

[52] Dupéré-Richer D, Licht JD. Epigenetic regulatory mutations and epigenetic therapy for multiple myeloma. Curr Opin Hematol 2017;24:336–44. 10.1097/MOH.0000000000000358.

[53] Zhang XY, Rajagopalan D, Chung T-H, Hooi L, Toh TB, Tian JS, et al. Frequent upregulation of G9a promotes RelB-dependent proliferation and survival in multiple myeloma. Exp Hematol Oncol 2020;9:8. 10.1186/s40164-020-00164-4.

[54] Ishiguro K, Kitajima H, Niinuma T, Maruyama R, Nishiyama N, Ohtani H, et al. Dual EZH2 and G9a inhibition suppresses multiple myeloma cell proliferation by regulating the interferon signal and IRF4-MYC axis. Cell Death Discov 2021;7:7. 10.1038/s41420-020-00400-0.

[55] Dunphy K, Bazou D, Henry M, Meleady P, Miettinen JJ, Heckman CA, et al. Proteomic and Metabolomic Analysis of Bone Marrow and Plasma from Patients with Extramedullary Multiple Myeloma Identifies Distinct Protein and Metabolite Signatures. Cancers 2023;15:3764. 10.3390/cancers15153764.

[56] Huang B, Liu H, Chan S, Liu J, Gu J, Chen M, et al. RUNX2 promotes the suppression of osteoblast function and enhancement of osteoclast activity by multiple myeloma cells. Med Oncol Northwood Lond Engl 2023;40:115. 10.1007/s12032-023-01960-8.

[57] Yang G, Hamadeh IS, Katz J, Riva A, Lakatos P, Balla B, et al. SIRT1/HERC4 Locus Associated With Bisphosphonate-Induced Osteonecrosis of the Jaw: An Exome-Wide Association Analysis. J Bone Miner Res Off J Am Soc Bone Miner Res 2018;33:91–8. 10.1002/jbmr.3285.

